# Identification of genomic alterations with clinical impact in canine splenic hemangiosarcoma

**DOI:** 10.1101/2022.11.17.516327

**Authors:** Timothy Estabrooks, Anastasia Gurinovich, Jodie Pietruska, Benjamin Lewis, Garrett Harvey, Gerald Post, Lindsay Lambert, Lucas Rodrigues, Michelle E. White, Christina Lopes, Cheryl A. London, Kate Megquier

## Abstract

**Background:** Canine hemangiosarcoma (HSA) is an aggressive cancer of endothelial cells associated with short survival times. Understanding the genomic landscape of HSA is critical to developing more effective therapeutic strategies.

**Objectives:** To determine the relationships between genomic and clinical features including treatment and outcome in canine splenic HSA.

**Animals:** 109 dogs with primary splenic HSA treated by splenectomy that had tumor sequencing via the FidoCure® Precision Medicine Platform targeted sequencing panel.

**Methods:** Patient signalment, weight, metastasis at diagnosis, treatment, and survival time were retrospectively evaluated. The incidence of genomic alterations in individual genes and their relationship to patient variables and outcome were assessed.

**Results:** Somatic mutations in *TP53* (n = 45), *NRAS* (n = 20), and *PIK3CA* (n = 19) were most common. Survival was associated with metastases at diagnosis, germline variants in *SETD2* and *NOTCH1*, and nominally with breed. Age at diagnosis was associated with *NRAS* mutations and breed. *TP53* and *PIK3CA* mutations were found in larger dogs, germline *SETD2* variants in smaller dogs. Doxorubicin (DOX) treatment did not significantly improve survival time, while targeted therapies had a significant early survival benefit.

**Conclusions and clinical importance:** DOX treatment may provide limited clinical benefit for dogs with splenic HSA, while targeted therapy may provide early survival benefit. Genetic signatures associated with splenic HSA may be useful in guiding targeted therapy to improve outcomes. Germline variants, age, size, and breed may be useful prognostic factors and provide insight into the genomic landscape of the tumor.

## 1 INTRODUCTION

Hemangiosarcoma (HSA) is a common, aggressive cancer in dogs that arises from endothelial progenitor cells, most frequently in the spleen.^1^ Despite aggressive treatment, median survival times range from 4-8 months due to a high metastatic rate and rapid tumor recurrence.^2^ Unfortunately, patient outcomes have not improved significantly in the past 30 years.^3,4^ In the era of precision medicine, understanding the genomic landscape of HSA will likely facilitate identification and implementation of new, more effective therapeutic strategies. This is particularly important given that canine HSA is a relevant large animal comparative model for human angiosarcoma (AS), a cancer that bears histologic and clinical similarities to canine HSA but occurs far less frequently (300-800 human cases/year compared to greater than 25,000 cases/year in dogs).^5,6^ As with canine HSA, human AS exhibits an aggressive biologic behavior including resistance to chemotherapy and the development of drug resistant metastasis, resulting in a 5-year survival rate of only 26%.^7^ Consequently, validating precision medicine approaches in canine HSA by leveraging its genomic landscape to guide therapy could provide critical new data regarding therapeutic combinations that may be of benefit for the human disease.

Previous studies have performed whole exome sequencing (WES) and targeted next-generation sequencing (NGS) of canine HSA tumors, finding potential driver mutations in *TP53, PIK3CA, NRAS*, and *PTEN*, among other genes.^8-12^ Loss-of-function mutations in the *TP53* tumor-suppressor gene were most frequent across studies (29-93% of cases), as well as activating mutations of *PIK3CA* (14-60%) and *NRAS* (4-24%).^8-12^ In one paper, increased PI3K pathway signaling was demonstrated in cases with either *PIK3CA* activating mutations or *PTEN* inactivating mutations, and increased MAPK/ERK pathway signaling was found in cases with *NRAS* activating mutations.^9^ While both AS and HSA are genetically heterogeneous, there are some similarities, including mutations in *TP53, PIK3CA* (most common in primary breast AS^13^), *PTEN*, and *NRAS*^14,15^, and MAPK/ERK and PI3K pathway activation.^14,15^

Sequencing of patient tumors is increasingly being used to identify targetable mutations and match patients to a “precision medicine” treatment. With mutation data and long-term follow-up, outcomes of such precision therapies have provided important information regarding efficacy of individual and combination treatment strategies for human cancer patients. In the current study, we leveraged targeted NGS data and matched clinical annotations in a population of 109 dogs with splenic HSA to assess associations between genomic features, clinical presentation, treatment regimens and outcome. This study represents the largest cohort of patients with splenic HSA to have undergone genomic interrogation, and the first to show a link between somatic variants and clinical variables such as age and weight, in addition to an association between germline variants, breed, and overall survival. Moreover, these data confirm prior published data questioning the efficacy of doxorubicin for the treatment of splenic HSA.^16^

## 2 MATERIALS AND METHODS

### 2.1 Case selection criteria

Cases were enrolled retrospectively by reviewing medical records of dogs with splenic HSA for which the splenic mass was submitted for NGS through the FidoCure Precision Medicine platform (One Health Company, Palo Alto, CA). Dogs were included if they had undergone splenectomy to remove the tumor, had a histologic diagnosis of HSA by a board certified veterinary anatomic pathologist, the splenic mass was presumed to be the primary tumor site, and tumor samples had been submitted for FidoCure NGS. Dogs were excluded if surgery to remove the splenic tumor was not performed or if the confirmed or suspected primary tumor was in a non-splenic location (*e*.*g*., right auricle, cutaneous, subcutaneous, retroperitoneal, *etc*.).

One hundred ten dogs with splenic HSA submitted for FidoCure analysis were identified. One dog was excluded because it could not be verified that the spleen was the primary tumor site, leaving 109 dogs for analysis. All but one dog had confirmed splenic HSA. The remaining dog had histopathology originally reported as splenic sarcoma and additional IHC to identify HSA was ordered by the requesting veterinarian. The mass was then submitted as HSA by the veterinarian, but the IHC results were not available for us to review so the diagnosis was presumed.

### 2.2 Data collection

Case records were retrospectively reviewed. Data collected included age at diagnosis, dog breed, sex, neuter status, weight, primary tumor site, presence and site of metastases, clinical stage, date of diagnosis, and date of death or last follow-up, and whether the dog received doxorubicin and/or targeted therapy. The presence and location of metastases at diagnosis was validated by review of patient medical records including imaging reports, histopathology of sampled metastatic sites, and the submitting veterinarian’s interpretation of in-house imaging. Presence of metastasis was determined by the submitting veterinarian via a variety of methods, including thoracic radiographs, abdominal ultrasound, thoracic and/or abdominal computed tomography (CT), and/or exploratory surgery. Metastasis was not confirmed by histopathology or cytology in all cases, but was often presumed based on imaging findings. Full staging with thoracic and abdominal imaging could not be confirmed in all cases, and in some cases the submitting veterinarian’s interpretation of imaging was available, but not the original imaging.

For purposes of treatment reporting, patients were considered to have been treated with doxorubicin (DOX) if they were reported to have received at least one dose of DOX at any point in their treatment. Targeted therapies were recommended by FidoCure based on the results of their NGS panel analysis of potentially targetable mutations. When recommended, these therapies could be ordered directly from FidoCure. Recommended therapies from FidoCure included rapamycin (an mTOR inhibitor), trametinib (a MEK inhibitor), vorinostat (an HDAC inhibitor), and multiple tyrosine kinase inhibitors; primarily toceranib, dasatinib, lapatinib, and imatinib. If patients ordered targeted therapies from FidoCure at any point, this was noted and the medication used was recorded. Total number of DOX doses received, targeted therapy doses received, and length of targeted therapy treatment could not be confirmed for all cases. It was not always possible to verify whether patients received other treatments beyond doxorubicin or FidoCure targeted therapies, such as other intravenous agents or metronomic chemotherapy.

### 2.3 Tumor sequencing

Sequencing of splenic HSA samples was performed using the NGS panel from the FidoCure Precision Medicine Platform targeting 56 individual genes (Supplementary Table 1). Tumor samples confirmed by histopathology were obtained as formalin-fixed paraffin-embedded (FFPE) tissues submitted by the clinic that had performed the splenectomy.

DNA was extracted from FFPE tissues using the Mag-Bind® FFPE DNA/RNA kit (Omega Bio-tek). DNA was quantified using the Qubit dsDNA HS assay kit (Thermo Fisher), and 200 ng was used to prepare a DNA sequencing library using the SureSelect Low Input Library (Agilent). Hybrid selection of the targeted regions was performed using the SureSelect custom DNA Target Enrichment Probes and SureSelect XT Hyb and Wash kit, following the manufacturer’s instructions. The final libraries were quantified using qPCR and pooled for sequencing. Samples were sequenced on Illumina MiSeq 2×150 or NovaSeq S4 2×150 sequencers to a target depth of approximately 500x. Sequencing reads were aligned to the CanFam3.1 reference genome^17^ using bwa-mem (v0.7.12).^18^ Preprocessing was performed using Picard Tools MarkDuplicates (http://broadinstitute.github.io/picard) and following the GATK^19^ (version 3.8.1) best practices. Bamtools^20^ was used to filter out reads with mapping quality less than five, or with ten or more mismatches. Sequencing metrics are provided in Supplementary Table 2. Much of this sequencing data was also included in a prior publication^21^ of a much larger set of tumor sequencing information for a diverse set of canine cancers. However, the data associated with splenic HSA did not undergo additional analysis for functional consequences of somatic mutations, recredentialling of the variant calls, and association with clinical outcomes.

Single-nucleotide variants (SNVs) and insertions and deletions (indels) were identified by creating a pileup file in SAMtools^22^ and calling variants using Varscan2^23^, requiring passing variants to have coverage > 10x, variant allele fraction (VAF) ≥1%, and minimum quality score of 20. Additional filtering was performed to remove variants with VAF < 2% or > 95% (potential sequencing artifacts or homozygous germline mutations), and variants located in repetitive regions^24^ were filtered out. Variants were phased using the tool WhatsHap^25^, and variants co-occurring in the same read (within 150 base pairs) were filtered out as putative germline variants. SnpEff and SnpSift^26^ were used to annotate each variant and predict its functional impact. Variants with moderate or high impact were included in downstream analysis.

Identified mutations were compared to known human mutations in the Catalog of Somatic Mutations in Cancer (COSMIC, cancer.sanger.ac.uk)^27^ to determine likelihood of being pathogenic. Due to the lack of a matched normal germline sample in variant calling, we could not definitively distinguish between all somatic mutations and germline variants. However, we annotated variants found in two catalogs of germline variants from 722 canids^28^ and from 591 dogs^29^, as well as common (≥ 5 cases) variants with VAF near 0.5 or 1 as putative germline variants in our cohort (Supplementary Table 3). Mutations identified in < 5 cases that did not overlap a known human pathogenic mutation were flagged as “unknown”. Variants remaining after filtering were annotated as somatic.

### 2.4 Statistical analysis and generation of figures

Overall survival time was defined as the time from HSA diagnosis to patient death or censoring. Patients were censored if they were lost to follow-up or still alive at the time of data analysis. All patient deaths were considered death from disease unless a clear, unrelated cause was confirmed. To identify any important clinical variables associated with OST, we used univariate linear regression models for continuous variables and one-way ANOVA tests for categorical variables. Survival function was estimated using the Kaplan-Meier method^30^ with median survival time (MST) and 95% confidence intervals and differences in overall survival time (OST) between groups evaluated using the log-rank and Wilcoxon signed-rank tests. A Cox Proportional Hazards model incorporating multiple clinical factors was also created. Factors evaluated for prognostic significance using Kaplan-Meier survival analysis included age at diagnosis, sex, weight, presence of metastasis at diagnosis, doxorubicin treatment, targeted therapy treatment, and total number of somatic mutations.

We also looked at the relationship between various clinical features and individual genes altered by somatic mutations or germline variants. We used linear regression models for continuous variables, such as OST, age at diagnosis, weight, and total number of somatic mutations; logistic regression models for categorical variables with two levels, such as presence of metastasis at diagnosis; and Fisher’s exact tests for categorical variables with more than two levels, such as breed and reproductive status. Cox Proportional Hazards models containing the group of somatic mutations or germline variants were also assessed.

All statistical analyses were performed using R software, version 4.1.2.^31^ Forest plots were created using the survival^32^ and survminer^33^ R packages. Oncoprint and mutual exclusivity analysis were done using CBioportal’s Oncoprinter.^34,35^ “Lollipop” plots of mutation positions were created using the tool Lollipops v1.5.2.^36^ Gene interaction plot was created using the maftools^37^ package for R.

## 3 RESULTS

One hundred nine dogs were included in the study. Characteristics and descriptive statistics of this cohort are listed in Table 1. The most common breed designation was mixed breed (n = 32), followed by golden retriever (n = 16), German shepherd (n = 14), and Labrador retriever (n = 14). Males were slightly overrepresented (62%), and most were neutered (93%), while all females were spayed. Complete individual patient demographic and gene mutation data are listed in Supplementary Table 3.

**Table 1.** Summary characteristics of the 109 dogs in the study, including age, weight, breed, stage, and number censored.

|  |  |  |
| --- | --- | --- |
| **Age (years)** | Median (range) | 10 (4-14) |
| **Sex** | MC | 59 |
|  | MI | 8 |
|  | FS | 42 |
| **Weight (kg)** | Median (range) | 29 (4.7-59.2) |
| **Breed** | Mixed breed | 32 |
|  | Golden retriever | 16 |
|  | German shepherd | 14 |
|  | Labrador retriever | 14 |
|  | Pit bull | 3 |
|  | Bichon Frise | 2 |
|  | Flat-coated retriever | 2 |
|  | Boxer | 2 |
|  | Australian shepherd | 2 |
|  | Other (1 each) | 23 |
| **Stage** | I/II | 67 |
|  | III | 33 |
|  | Unknown | 9 |
| **# Censored** | Alive | 4 |
|  | Lost to follow-up | 9 |

### 3.1 Clinical variables

We examined the relationship between clinical variables, including age at diagnosis, sex and neuter status, weight, presence of metastasis at diagnosis, and treatment with overall survival (OST). Univariate and multivariate regression analyses were performed, along with cox proportional hazards models to identify important factors (Figure 1).

**Figure 1.**
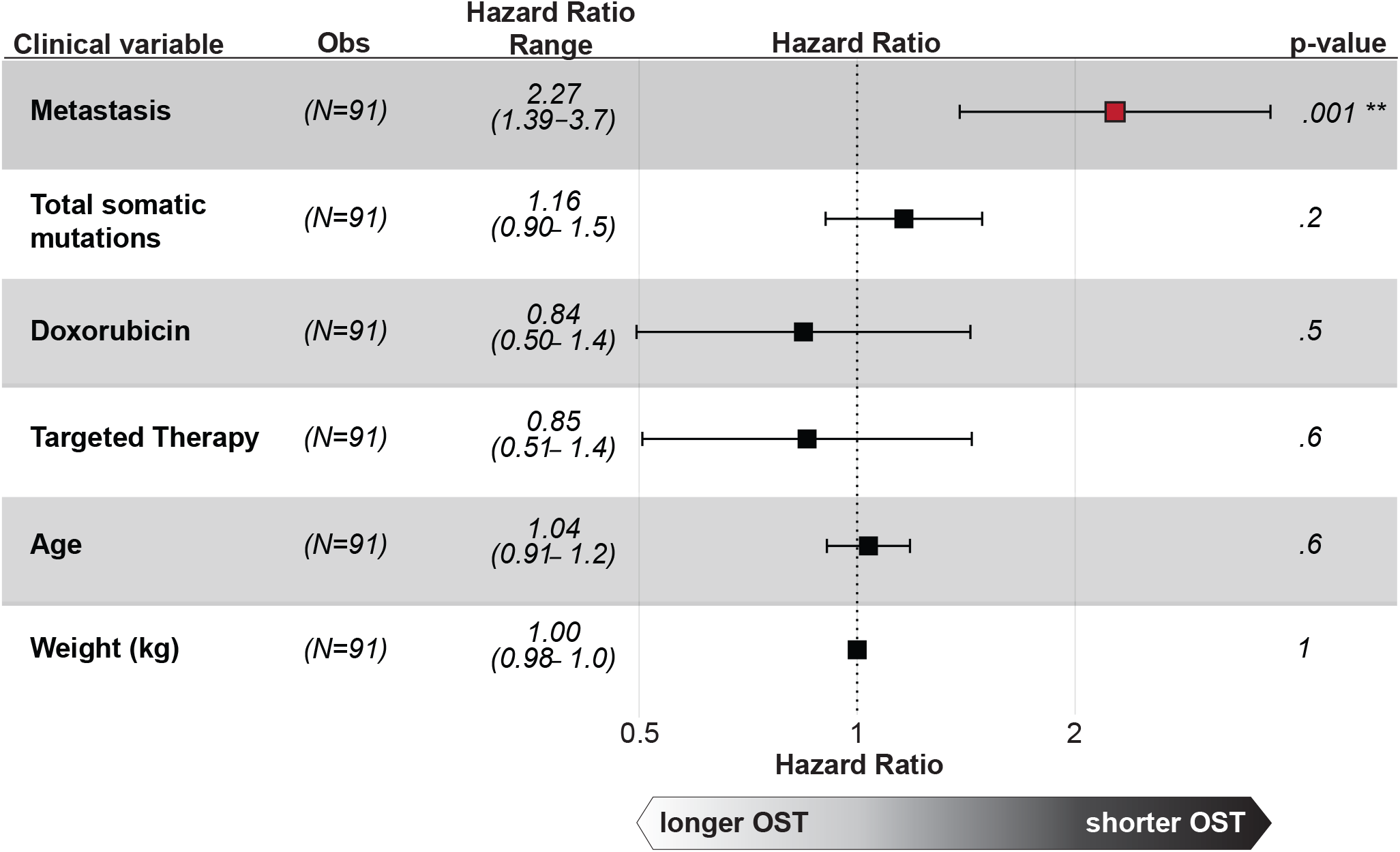
Effect of clinical variables on survival. Forest plot of Cox Proportional Hazards model effects of multiple clinical variables on overall survival (OST). Dogs with detectable metastases at the time of diagnosis had a significantly shorter overall survival.

#### Progression free and overall survival

Complete treatment information was available for 99 dogs. Outcome was available for 100 dogs, with nine lost to follow-up or still alive at the time of data collection. The MST of all patients was 166 days (range, 16-956 days) (Supplementary Figure 1A). Ninety-six dogs were dead of disease at the time of data collection, while four were still alive and nine were lost to follow-up. Age at diagnosis, weight, total number of somatic mutations, and sex/neuter status had no significant effect on OST (Figure 1).

Thirty-three dogs had confirmed or presumed metastasis, with the liver (n = 25) and omentum (n = 4) being the most frequent sites of metastasis. Nine dogs could not be classified because data regarding metastasis at diagnosis was not available. Dogs with metastasis at diagnosis had significantly shorter survival compared to those without metastasis (*P =* <.001), with an MST of 120 days (range [95% CI], 16-596 days [87-156 days]) versus 252 days (range [95% CI], 36-956 days [207-365 days]) respectively (Figure 2A).

**Figure 2.**
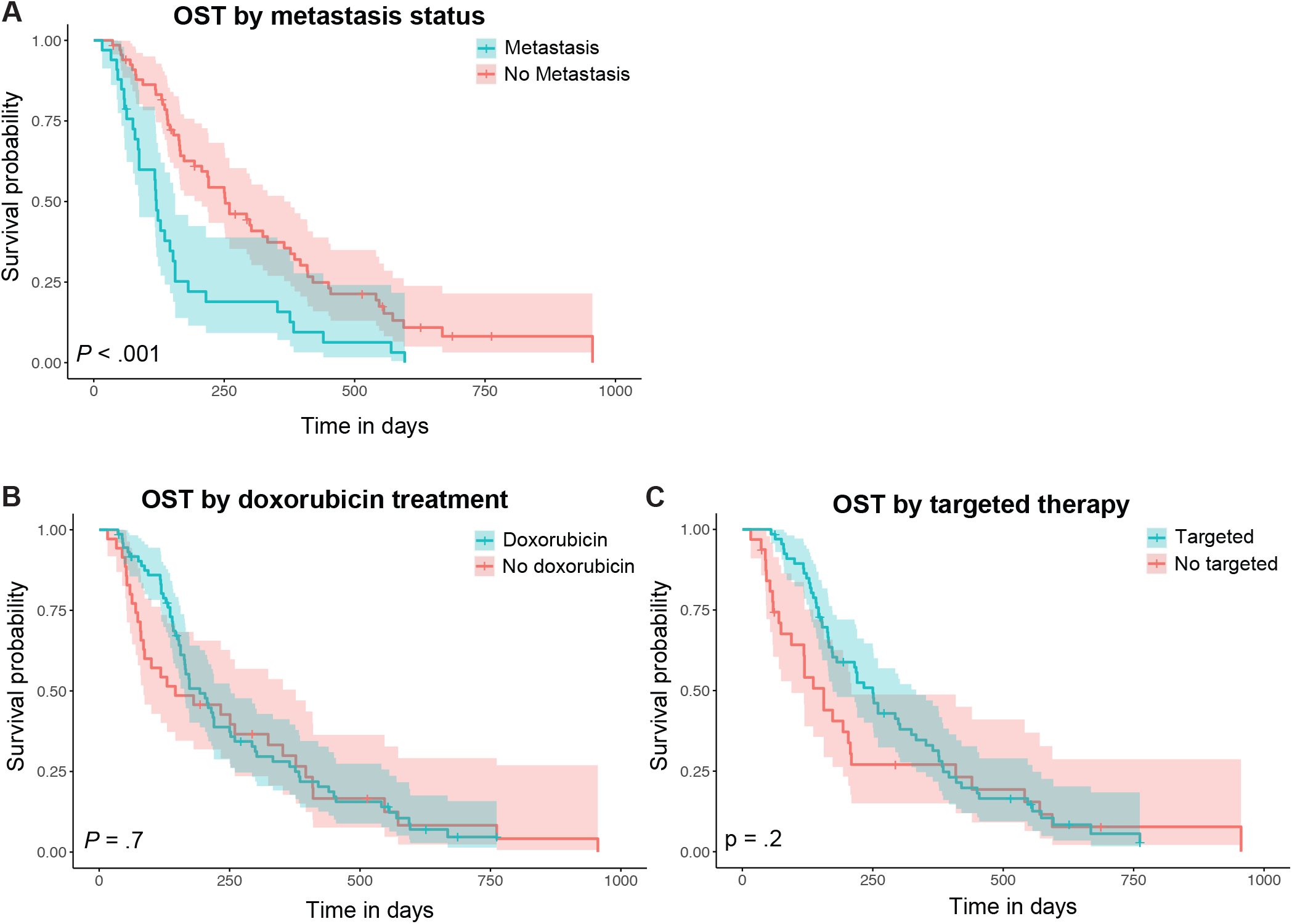
Clinical variable survival curves. Kaplan-Meier survival curves comparing dogs (A) with or without metastases at time of presentation; (B) with and without doxorubicin therapy; (C) with and without targeted therapy.

**Figure 3.**
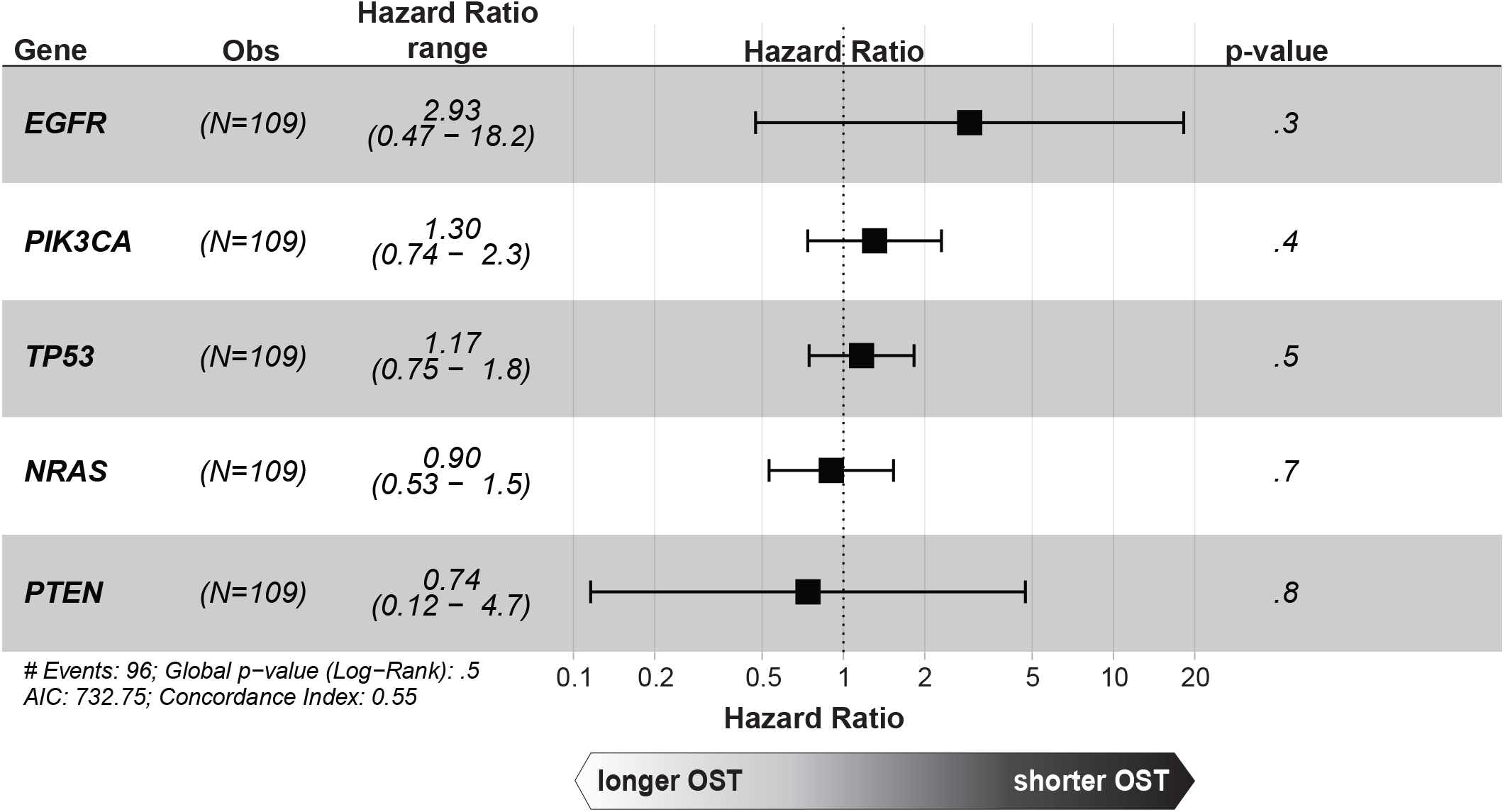
Effect of somatic mutations on survival. Forest plot of Cox Proportional Hazards model effects of the most commonly somatically mutated genes on OST. None had a significant effect.

Following splenectomy, 73 dogs received at least one dose of DOX and 67 dogs ordered the FidoCure-recommended targeted therapy. Of the 67 dogs that received targeted therapy, 45 also received DOX, although relative timing of the two therapies is unknown. Ten dogs were missing data with respect to the type of treatment given; one dog had an unknown DOX treatment status and treatment with FidoCure-recommended therapy could not be confirmed in all 10. Of the 67 dogs that ordered targeted therapies, 42 ordered more than one medication. Twenty-four dogs received only DOX, while 22 dogs received only targeted therapy. Eight dogs did not receive either DOX or targeted therapy.

Treatment with DOX did not significantly improve survival time in the combined population (*P* =.7). Dogs that received at least one dose of doxorubicin had an MST of 193 days (range [95% CI], 36 - 762 days [163 - 252 days]) while dogs that did not receive doxorubicin had an MST of 146 days (range [95% CI], 16 - 956 days [85 - 377 days]) (Figure 2B). Early survival subjectively appeared to be improved with doxorubicin chemotherapy, but this was not statistically significant (Wilcoxon test *P =*.2).

Patients that received targeted therapy (plus or minus doxorubicin treatment) had a longer survival time compared to those that did not (MST of 250 days (range [95% CI], 55 - 762 days [173-333 days]) versus 156 days (range [95% CI], 16 - 956 days [94-209 days]). This difference was significant on linear regression (*P*_*Wilcoxon*_ =.003, *P*_*adj*_ =.007), with early survival improved, but was not significant for overall survival by Kaplan Meier (*P*_*KM*_ =.2) or Cox Proportional Hazards model (*P*_*Cox*_ =.6) (Figure 2C).

### 3.2 Genomic landscape

Sequenced DNA libraries achieved a mean depth of 1704x overall (range, 74-8661). On average, 99.7% of reads aligned to the canine genome (range, 97% - 100%), with an average duplicate read percentage of 19% (range, 0% - 38%) (Supplementary Table 1).

After variant calling and filtration, somatic mutations were identified in 72 cases, germline variants in 99 cases, and both in 65 cases (Supplementary Figure 2, Supplementary Table 3). Two cases had no detected mutations in the 56 genes targeted by the Fidocure NGS panel, and one case had no mutations remaining after filtering (Supplementary Table 1). The mean number of somatic mutations per case, including cases with multiple mutations in the same gene, was 1.1 (range, 0 - 4).

#### Somatic mutations

Three genes (*TP53, NRAS*, and *PIK3CA*) were somatically mutated in at least ten cases (Table 2). *TP53 (n =* 45 cases) was most commonly altered, with the majority of mutations present in the DNA-binding domain (Supplementary Figures 2-3, Supplementary Table 3). *NRAS* and *PIK3CA* were mutated in 20 and 19 cases, respectively. Somatic mutations in *PTEN* and *EGFR* were identified in only 3 cases, but were included in the overall analysis due to prior published data demonstrating their importance in canine HSA. Somatic mutations in three genes were associated with higher numbers of total somatic mutations *TP53* (*P*_*adj*_ <.001), *PIK3CA* (*P*_*adj*_ = <.001), and *PTEN* (*P*_*adj*_ =.006).

**Table 2.** Summary of genes with somatic mutations in 10 or more cases, including total number of individual mutations per gene.

| Gene | Cases | Individual Mutations |
| --- | --- | --- |
| *TP53* | 45 | 55 |
| *NRAS* | 20 | 20 |
| *PIK3CA* | 19 | 19 |

#### Germline variants

Putative germline variants were identified in ten or more cases in 11 genes, including *NOTCH1* (29 cases), *ROS1* (29 cases), *KMT2C* (26 cases), and *MET* (22 cases) (Table 3, Supplementary Table 3). Cases with germline *EGFR* variants were also included, despite falling below the cutoff (n = 6).

**Table 3.** Summary of genes with germline variants in 10 or more cases, including total number of individual variants per gene.

| Gene | Cases | Individual Variants |
| --- | --- | --- |
| *NOTCH1* | 32 | 38 |
| *ROS1* | 30 | 32 |
| *KMT2C* | 28 | 32 |
| *KMT2D* | 25 | 32 |
| *MET* | 22 | 22 |
| *BRCA2* | 21 | 26 |
| *PDGFRB* | 21 | 23 |
| *CDKN2A* | 17 | 18 |
| *BRCA1* | 14 | 37 |
| *FLT3* | 14 | 15 |
| *PARP1* | 12 | 13 |
| *SETD2* | 11 | 13 |

#### Co-occurrence and mutual exclusivity of somatic and germline alterations

We noted patterns of co-occurrence and mutual exclusivity in both somatic mutations and germline variants (Supplementary Table 4, Supplementary Figure 4). Somatic mutations in six pairs of genes had nominally significant patterns of co-occurrence or mutual exclusivity that were not significant after multiple testing correction. *TP53* and *PIK3CA* mutations (co-occuring, *P* =.004, *P*_*adj*_ =.06), *TP53* and *NRAS* (mutually exclusive, *P* =.01, *P*_*adj*_ =.14), *PIK3CA* and *NRAS* (mutually exclusive, *P* = 0.02, *P*_*adj*_ =.2). Similarly, germline variants in *KMT2C* and *CDKN2A* (*P* =.02, *P*_*adj*_ =.2) and *PDGFRB* and *NOTCH1* (*P* =.03, *P*_*adj*_ =.3), mutations tended to be mutually exclusive although again these were not significant after multiple testing correction. Both somatic and germline alterations in *EGFR* nominally tended to co-occur with germline variants in *BRCA2* (*P* =.05, *P*_*adj*_ =.4).

#### Survival

None of the somatically mutated genes were associated with OST. However, germline variants in *SETD2* were associated with decreased OST in a Cox Proportional Hazards model (*P* =.001, Figure 4) and Kaplan-Meier survival analysis (*P*_*KM*_ <.001, Figure 5), with an MST for mutated and non-mutated cases of 84 days (range [95% CI], 45 - 250 days [59 days – NA]) and 207 days (range [95% CI], 16 - 956 days [165-260 days]), respectively. Germline variants in *NOTCH1* were also associated with decreased survival in a Cox Proportional Hazards model (*P*_*cox*_ =.04, Figure 4) and Kaplan-Meier survival analysis (*P*_*KM*_ =.04, MST for mutated and non-mutated cases of 165 days (range [95% CI], 33 – 556 days [146 – 260 days]) and 203 days (range [95% CI], 16 - 956 days [146 - 324 days]), respectively (Figure 5).

**Figure 4.**
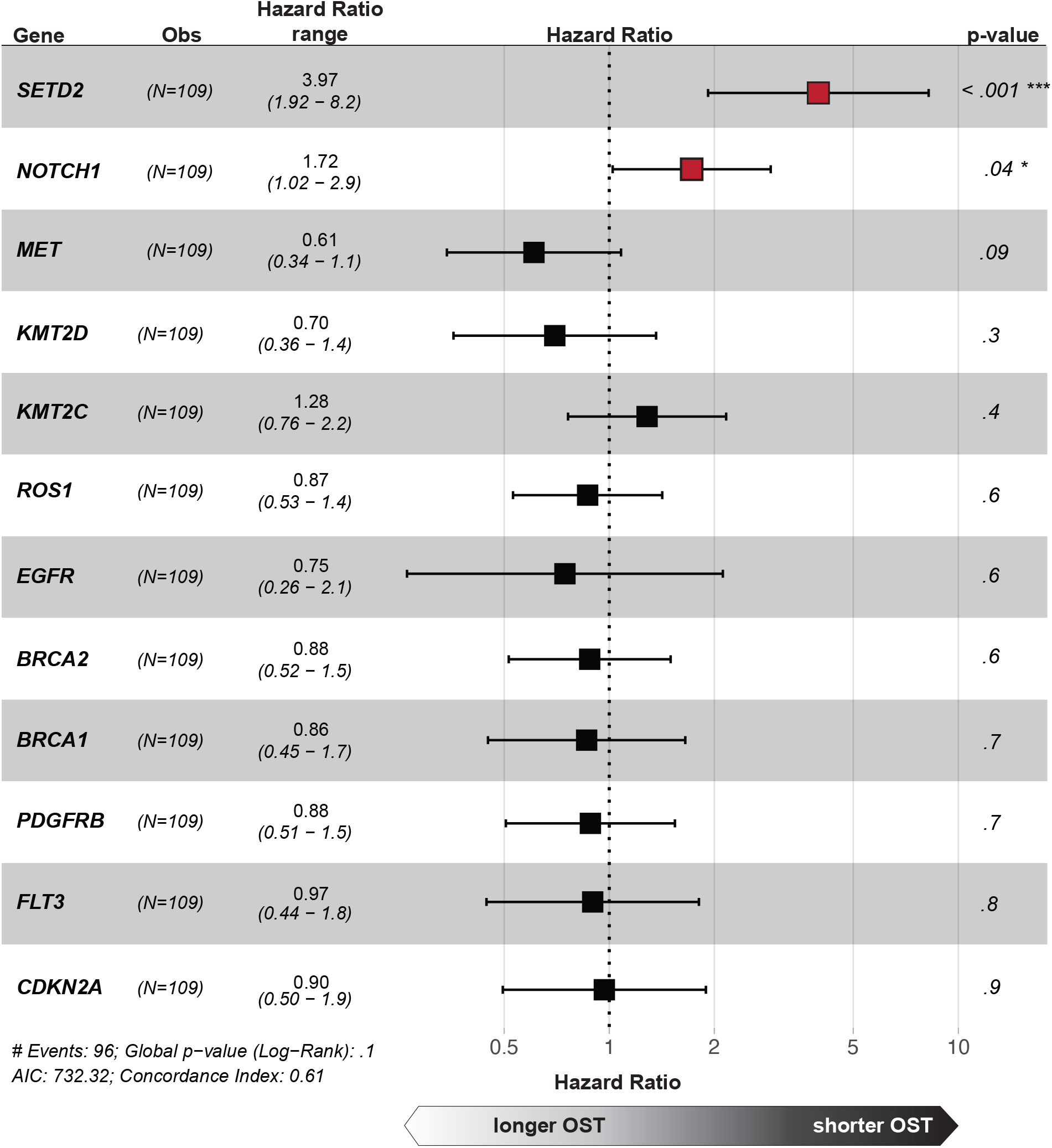
Effect of germline variants on survival. Forest plot of Cox Proportional Hazards model effects of the genes most commonly harboring germline variants on OST. Variants in *SETD2* and *NOTCH1* both were associated with significantly shorter OST.

**Figure 5.**
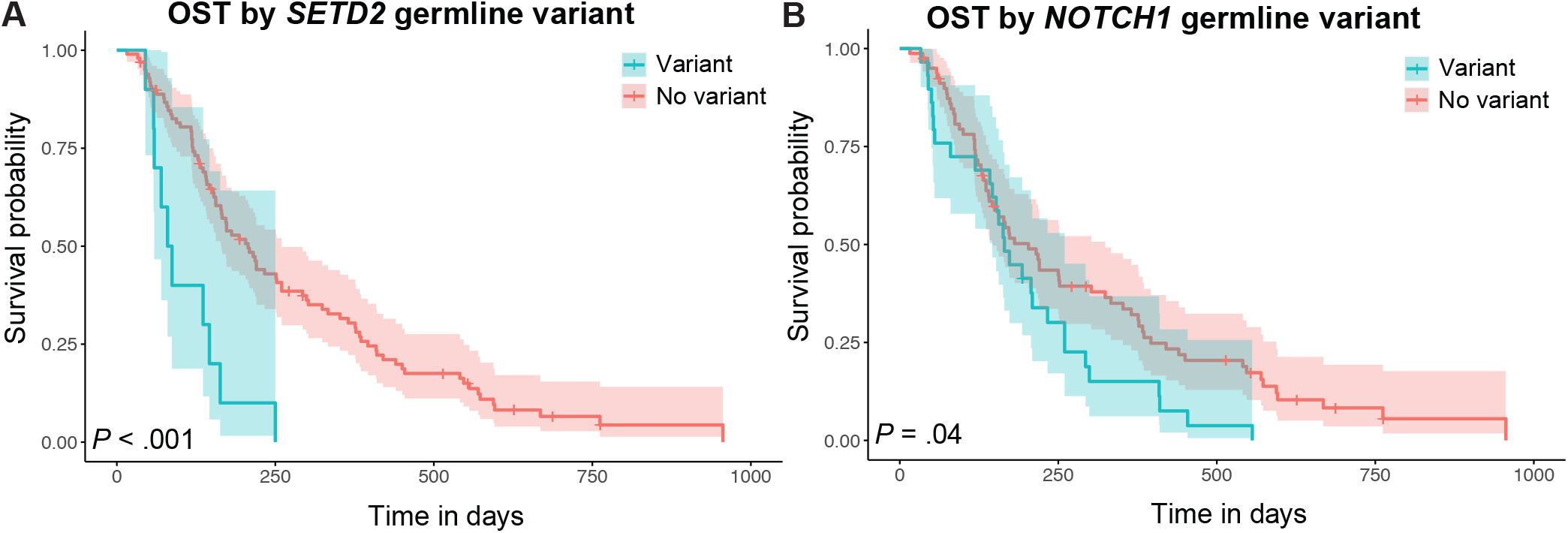
Germline variant survival curves. Kaplan-Meier curves comparing the survival of dogs with and without germline variants in (A) *SETD2*; and (B) *NOTCH1*.

#### Age

Age at diagnosis was available for all dogs (mean, 9.7 years; range, 4 - 14 years) (Supplementary Table 3, Supplementary Figure 1B). The association between age at diagnosis and individual genes was evaluated using a univariate linear regression model. Somatic mutations in *NRAS* were significantly associated with younger age of diagnosis (unadjusted *P* <.001; *P*_*adj*_ =.004), with a mean age of dogs carrying *NRAS* mutations 8 years (range, 4 - 13 years) vs. 10 years (range, 6 - 14 years) in those without *NRAS* mutations (Supplementary Figure 5). No other genes with somatic mutations were associated with age.

Germline variants in *KMT2C* and *SETD2* were nominally associated with increased age, but this was not significant after correction for multiple testing. Cases with *KMT2C* variants had a mean age of 11 years (range, 7 - 14) vs. 10 years in cases without (range, 4 - 14) (*P* =.007; *P*_*adj*_ =.08). Cases with *SETD2* variants had a mean age of 10.6 years (range, 7 - 14) vs. 9.4 years in cases without a variant (range, 4 - 14) (*P* =.04; *P*_*adj*_ =.3)(Supplementary Figure 5).

#### Weight

Weights were available for all 109 dogs (mean = 29.0 kg, range = 4.7 - 59.2) (Supplementary Figure 1C). On univariate linear regression, mean weights were significantly different for dogs with somatic mutations in *PIK3CA* (*P*_*adj*_ *=*.03) and *TP53* (*P*_*adj*_ *=*.03). In both cases, dogs carrying mutations tended to be larger than dogs without somatic mutations (*PIK3CA*: mean with mutation = 35.2 kg, mean without mutation = 27.7 kg; *TP53*: mean with mutation = 32.3 kg, mean without mutation = 26.7 (Supplementary Figure 6).

Dogs carrying germline variants in *SETD2* tended to be smaller than those without (mean weight with variant = 20 kg, mean weight without variant = 29.9 kg, *P* =.007), however this was not significant after multiple testing correction(*P*_*adj*_ =.09) (Figure 5).

#### Breed

There were three breeds for which sample numbers were sufficient to evaluate the distribution of various variables: golden retrievers (n = 16), German shepherd dogs (n = 14), and Labrador retrievers (n = 14). The mean age at diagnosis differed significantly among these three breeds (*P* =.01, one way ANOVA test), with German shepherd dogs (mean age = 8.5 years) being younger than golden retrievers (mean age = 9.4 years) and Labrador retrievers (mean age = 10.6 years) being older (Figure 6). A nominal difference in OST was also noted, with German shepherd dogs having a shorter OST (median, 138 days; range, 16 - 514) and Labrador retrievers having a longer OST (median, 340 days; range, 36 - 594) than golden retrievers (median, 179 days; range, 87 - 596), but this was not statistically significant (*P* =.07)(Figure 6).

**Figure 6.**
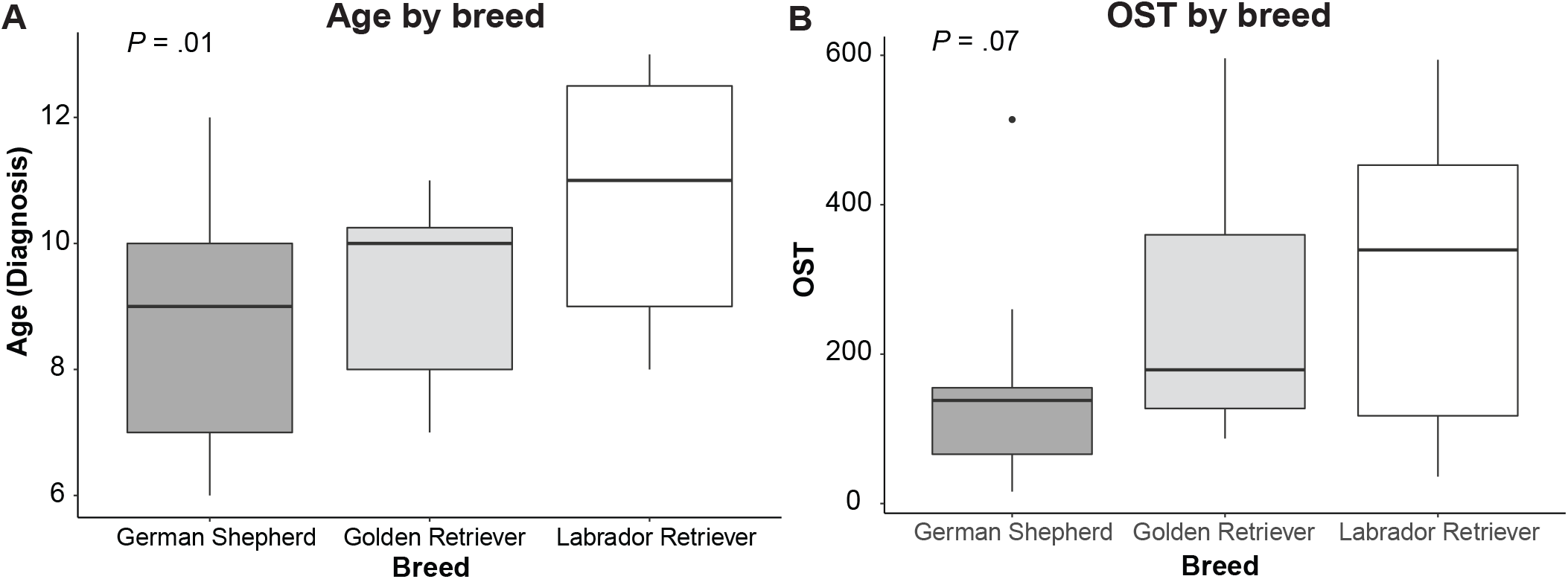
Breed effects. Box plots comparing (A) age and (B) OST in the three most common breeds in the cohort, golden retrievers (n = 16), German shepherd dogs (n = 14) and Labrador retrievers (n = 14).

A significant association with breed was also found for germline variants in *CDKN2A*, with only German shepherds among the three breeds included in the analysis carrying variants in this gene (*P* <.001; *P*_adj_=.009, Supplementary Figure 7B). Nominally significant differences in breed distribution that were not significant after correcting for multiple testing were noted in *TP53* (*P* =.02; *P*_*adj*_ =.08), *ROS1* (*P* =.02; *P*_*adj*_=.07), *FLT3* (*P* =.02; *P*_*adj*_=.07), and *PDGFRB* (*P* =.03; *P*_*adj*_=.09)(Supplementary Figure 7B-E).

## 4 DISCUSSION

This study represents the largest exome sequencing study of primary canine splenic HSA to date, the first to link somatic mutations to patient characteristics such as age and size, and the first to link germline variants to patient outcome. Analysis of this data set confirms previously published data questioning the impact of doxorubicin on patient survival.^16^ Moreover, our findings suggest that treatment with targeted therapies may improve early survival.

### Clinical variables

In general, survival times were observed to be longer in this study than those previously reported. Non-treated patients had a median OST of approximately 5 months, while prior publications have reported survival times of 2-3 months for this population.^16,38,39^ Indeed, the dogs that received no treatment had a median OST comparable to a recent study comparing outcomes of dogs with HSA given carboplatin versus doxorubicin post-surgery (160 days and 139 days, respectively).^4^ This difference in OST may be influenced by immortal time bias^40^, as an unknown number of cases with more aggressive disease may have died before having the opportunity to enroll (or complete enrollment) in FidoCure.

The survival benefit of DOX in this population appeared to be primarily during the early course of treatment, with an improved surviving fraction observed in the first 3-4 months after diagnosis, although this was not statistically significant. Our data supports a prior study, in which dogs with HSA that received DOX post-surgery did not have a significantly improved survival time across the entire follow-up period compared to those that did not receive doxorubicin, however a significantly higher proportion of treated patients did survive within the first 4 months after surgery.^16^

Patients that received targeted therapies recommended by FidoCure had a longer OST versus those that did not (MST 250 days vs. 156 days), which was statistically significant upon univariate linear regression analysis. When Kaplan-Meier analysis was employed, this difference was statistically significant only during the early period of treatment, as the survival curves for treated and untreated patients reached equivalence at 375 days. Future prospective studies will be necessary to both identify targeted therapies best matched to the tumor genomic landscape, and to confirm benefit in the setting of single or multi-agent targeted therapy.

### Somatic mutations

Common somatic mutations in this cohort - *TP53* (41%, 30% - 93% previously reported), *NRAS* (18%, 0% - 24% previously reported), *PIK3CA* (17%, 15% - 60% previously reported), and *PTEN* (3%, 0% - 10% previously reported) were present at similar frequencies to previous reports. ^8-12^ However, mutations in both *TP53* and *PIK3CA* were observed near the lower end of their reported frequency in the literature. Our finding that *TP53* and *PIK3CA* are mutated more frequently in larger dogs may offer a potential explanation of the decreased prevalence of these mutations in our cohort, which included dogs as small as 4 kg.

We observed co-occurrence or mutually exclusive patterns of certain somatic mutations and germline variants, suggesting possible overlap in downstream effects. *PIK3CA, PTEN*, and *NRAS* mutations were mutually exclusive. *TP53* mutations frequently co-occurred with *PIK3CA/PTEN* mutations, while being mutually exclusive with *NRAS* mutations. Mutations in *PIK3CA* and *PTEN* are expected to have similar consequences, as PIK3CA promotes signaling through the PI3K/AKT/mTOR pathway and PTEN is a negative regulator of this signaling.^41-43^ *NRAS* mutations activate the RAS/RAF/MEK/ERK pathway, which has also been implicated in neoplastic development^14^, but these have also been shown to activate PI3K signaling as a downstream effector.^44,45^ Furthermore, evidence suggests that knockdown of TP53 can activate RAF/MEK/ERK independent of RAS, while RAS activation can inhibit TP53-mediated cell-cycle arrest.^46^ These data suggest the possibility that tumors with *PIK3CA/PTEN* mutations plus *TP53* mutation and tumors with *NRAS* mutations may be achieving similar downstream effects of RAF/MEK/ERK and PI3K activation and TP53 inhibition.

Overall, the patterns of co-occurrence/mutual exclusivity of both somatic mutations and germline variants suggest that key pathway aberrations driving disease pathogenesis can be achieved through germline or somatic genetic alteration of particular combinations of genes. Consequently, a more global view of both somatic and germline changes could be informative both for prognostication and therapy selection.

### Mutational burden

Our finding that overall mutational burden is correlated with somatic mutations in *TP53* replicates prior published work.^47^ *PIK3CA* and *PTEN* have not previously been linked to higher mutational burden, however, in this study, mutations in both genes tend to co-occur with *TP53* mutations, potentially confounding the analysis.

### Germline background

Due to the high prevalence of cancers within specific dog breeds, it is thought that many breeds carry fixed or high-frequency deleterious variants predisposing to cancer. Presumptive germline loci associated with HSA risk have been reported for golden retrievers.^48^ Many of the common germline alterations in this study fall into the RTK-RAS pathway, upstream of the MAPK pathway. The genomic profiles of canine HSA samples were previously found to have less enrichment in the MAPK pathway than human AS, suggesting a possible role of the germline background in altering MAPK signaling. Our findings highlight the potential role of germline background in the development of HSA in different breeds, and, importantly, in outcome. Variants in *SETD2* and *NOTCH1* were associated with decreased OST. *SETD2* is a known tumor suppressor gene encoding a histone methyltransferase, and decreased SETD2 protein expression or loss of function mutations have been implicated in tumor progression and poor prognosis in multiple human cancers, including gastric, pancreatic, lung, and renal cancers.^49-52^ Mutations have also been recently identified in canine osteosarcoma.^53,54^

The significant differences in age and suggestive differences in OST we observed between German Shepherds (which were younger and had shorter survival times), golden retrievers, and Labrador retrievers (which were older and had longer survival times) also point to the potential influence of the genetic background of these breeds. As germline *CDKN2A* variants were significantly more likely to occur in German Shepherds, it is possible that these variants contribute to underlying risk in this breed.

*CDKN2A* is a known tumor-suppressor gene and frequent deletions and copy number losses have been documented in both radiation-induced AS and canine HSA.^55^ Because these three breeds have different average lifespans, it is difficult to ascertain whether German Shepherds (mean lifespan of 10.3 years) age more rapidly compared to Golden retrievers (mean lifespan 12 years) and Labrador retrievers (12.6 years) ^56-58^, or if instead they age at the same rate, but their strong predisposition to HSA^56-59^ and tendency towards shorter OST depress the breed’s overall average survival. As golden retrievers and Labrador retrievers also have a high HSA risk, differences in genetic background and aging may be playing a role.

### Limitations

In addition to the previously stated limitations of this study’s retrospective design, and the difficulty of interpreting OST due to immortal time bias, there were also limitations in the sequencing methods. Patient tumor samples were sequenced without matched normal tissues, making it impossible to definitively distinguish germline mutations from somatic mutations. In addition, the targeted panel of 56 genes is fairly small, and we are unable to evaluate potential drivers not included in the panel, copy number changes, or other structural variants. In addition, despite the large cohort size, we did not have power to evaluate the efficacy of individual targeted therapies or combinations of therapies.

In conclusion, this study contributes significantly to our knowledge of the genomic landscape of primary canine splenic HSA, including impact of age, size, breed and genetic background may influence clinical presentation and outcome in this disease. Our findings also support the notion that the previously established standard of care cytotoxic chemotherapy (DOX) may not impact patient outcomes, providing a solid rationale for further research regarding the benefits of precision medicine in the setting of HSA. Prospective work to refine matching of genomic landscapes with appropriate targeted therapies in dogs may also facilitate improving outcomes of humans with AS.

## Abbreviations

AS: angiosarcoma
CI: confidence interval
CT: computed tomography
HSA: hemangiosarcoma
MST: median survival time
NGS: next-generation sequencing
OST: overall survival time
WES: whole exome sequencing

## Acknowledgments

This study was partially supported by a grant from the PETCO Foundation. The authors acknowledge the support of the One Health Company in supplying patient data and writing of the materials and methods.

