## Supplementary Figures for "Identification of genomic alterations with clinical impact in canine splenic hemangiosarcoma"

**Supplementary Figure 1**
**Distribution of clinical variables in cohort.** Histograms showing the count of dogs in each range bin for (A) overall survival time in days; (B) age in years; (C) weight in kilograms; and (D) total number of somatic mutations.

**Supplementary Figure 2**
**Oncoprint.** Oncoprint with each case in the cohort depicted as a column, showing clincial annotations (top); somatic mutations in five genes (middle); and germline variants in 12 genes (bottom).

**Supplementary Figure 3**
**Lollipop plots.** Lollipop plots showing the position of either somatic mutations or germline variants in selected proteins. Uniprot ID used to create each plot noted in lower right.

**Supplementary Figure 4**
**(A) Gene alteration interaction matrix.** Co-occurrence and mutual exclusivity of selected somatic mutations and germline variants. **(B) Heatmap of similarity scores.** Heatmap depicting the simple matching coefficients (SMC) between pairs of selected somatic mutations or germline variants.

**Supplementary Figure 5**
**Genes associated with age at diagnosis.** Boxplots depicting age at diagnosis in cases with or without (A) somatic mutations in *NRAS*; (B) germline variants in *KMT2C*; (C) germline variants in *SETD2*.

**Supplementary Figure 6**
**Genes associated with body weight.** Boxplots depicting weight in kilograms (kg) in cases with or without (A) somatic mutations in *TP53*; (B) somatic mutations in *PIK3CA*; (C) germline variants in *SETD2*.

**Supplementary Figure 7**
**Distribution of gene alterations by breed.** Bar charts depicting distribution in the top three most common breeds in the cohort of gene alterations in: (A) *TP53*; (B) *CDKN2A*; (C) *ROS1*; (D) *FLT3*; (E) *PDGFRB*.

#### Supplementary Figure 1

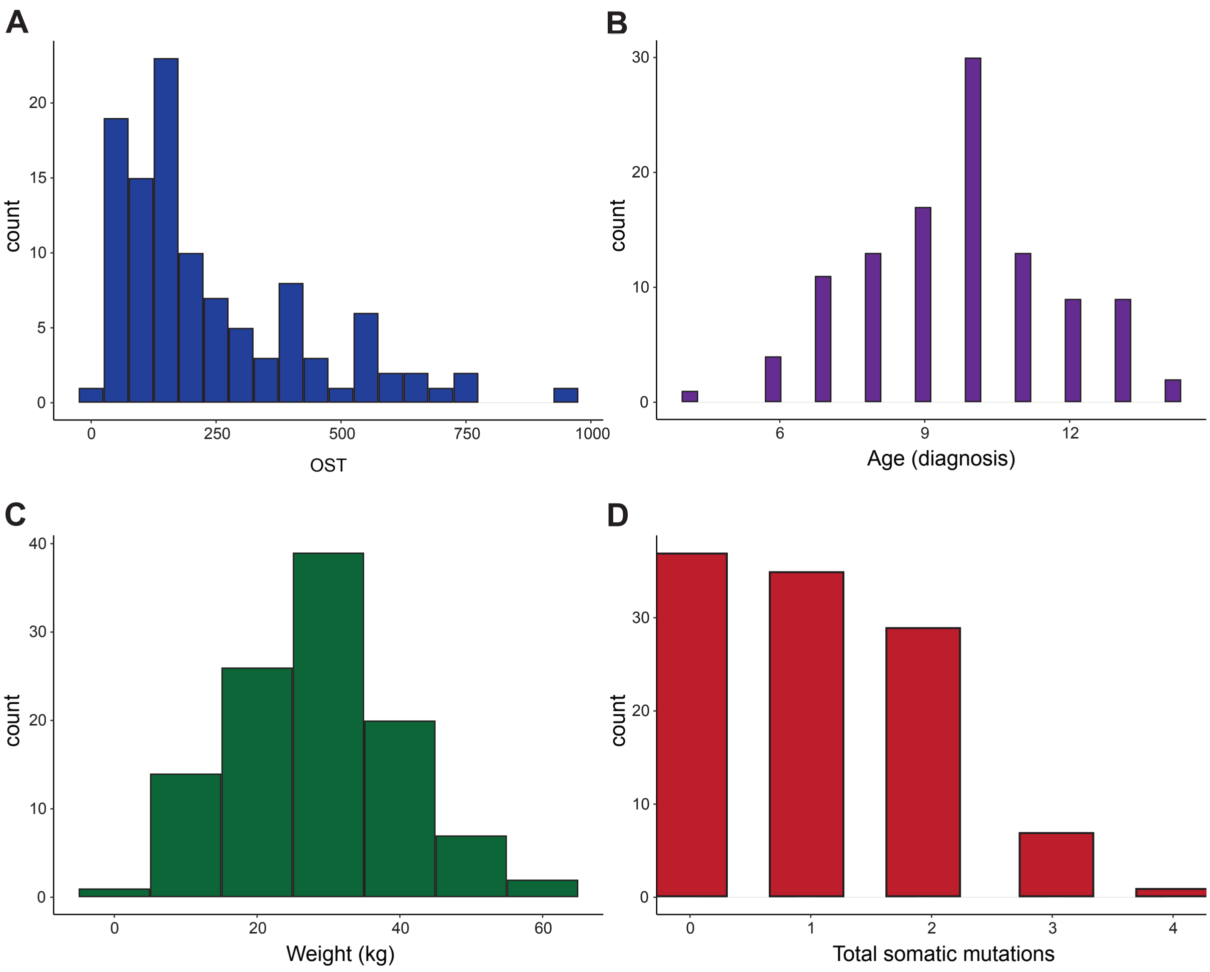

#### Supplementary Figure 2

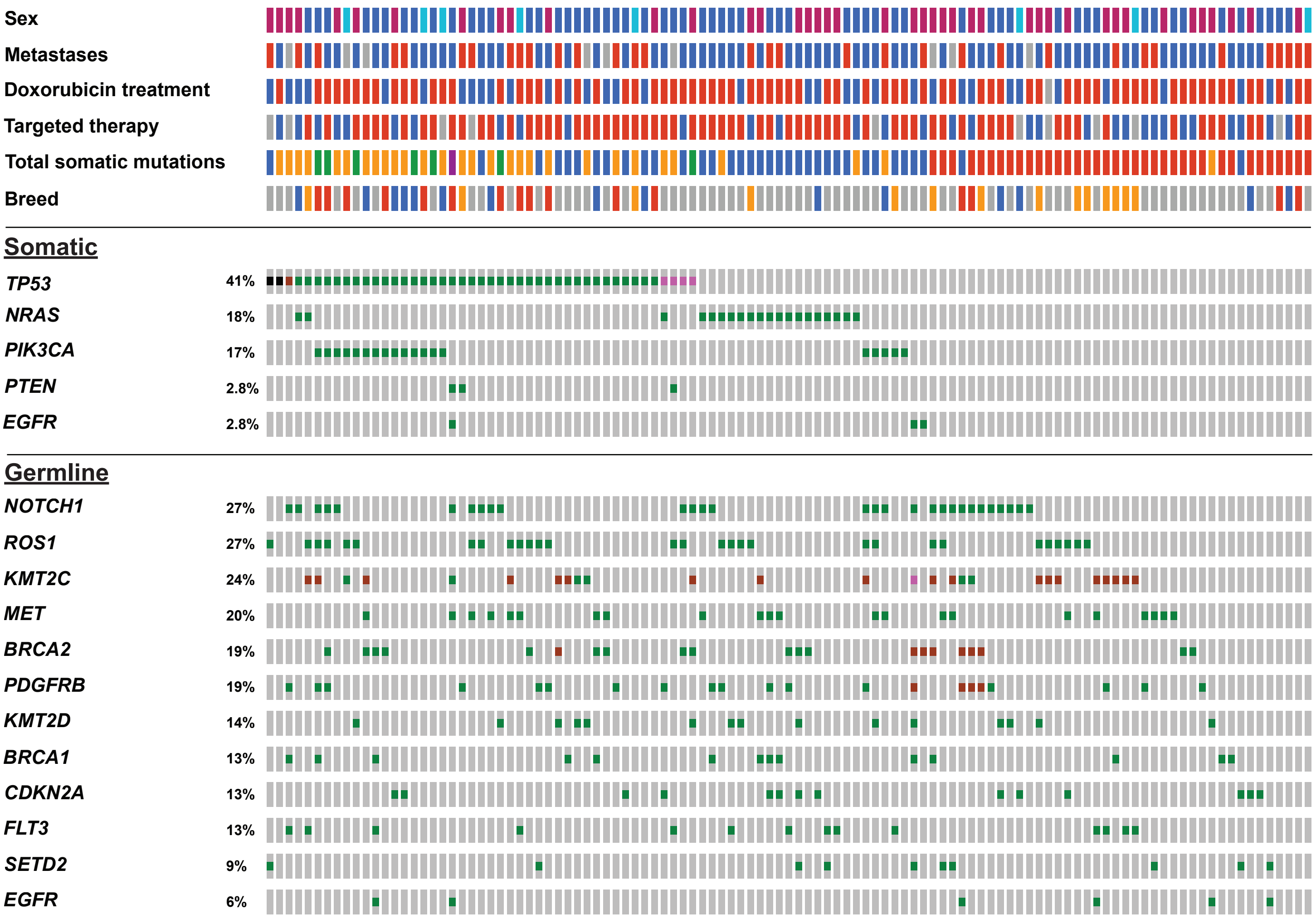


| | | | | | |
|---|---|---|---|---|---|
| **Genetic Alteration** | Inframe | Missense | Frameshift | Truncating | No alterations |
| **Sex** | FS | MI | MN | | |
| **Metastasis<br>Doxorubicin<br>Targeted therapy** | No | Yes | Unknown | | |
| **Breed** | golden retriever | Labrador retriever | German shepherd | Other | |
| **Total somatic mutations** | 0 | 1 | 2 | 3 | 4 |

#### Supplementary Figure 3

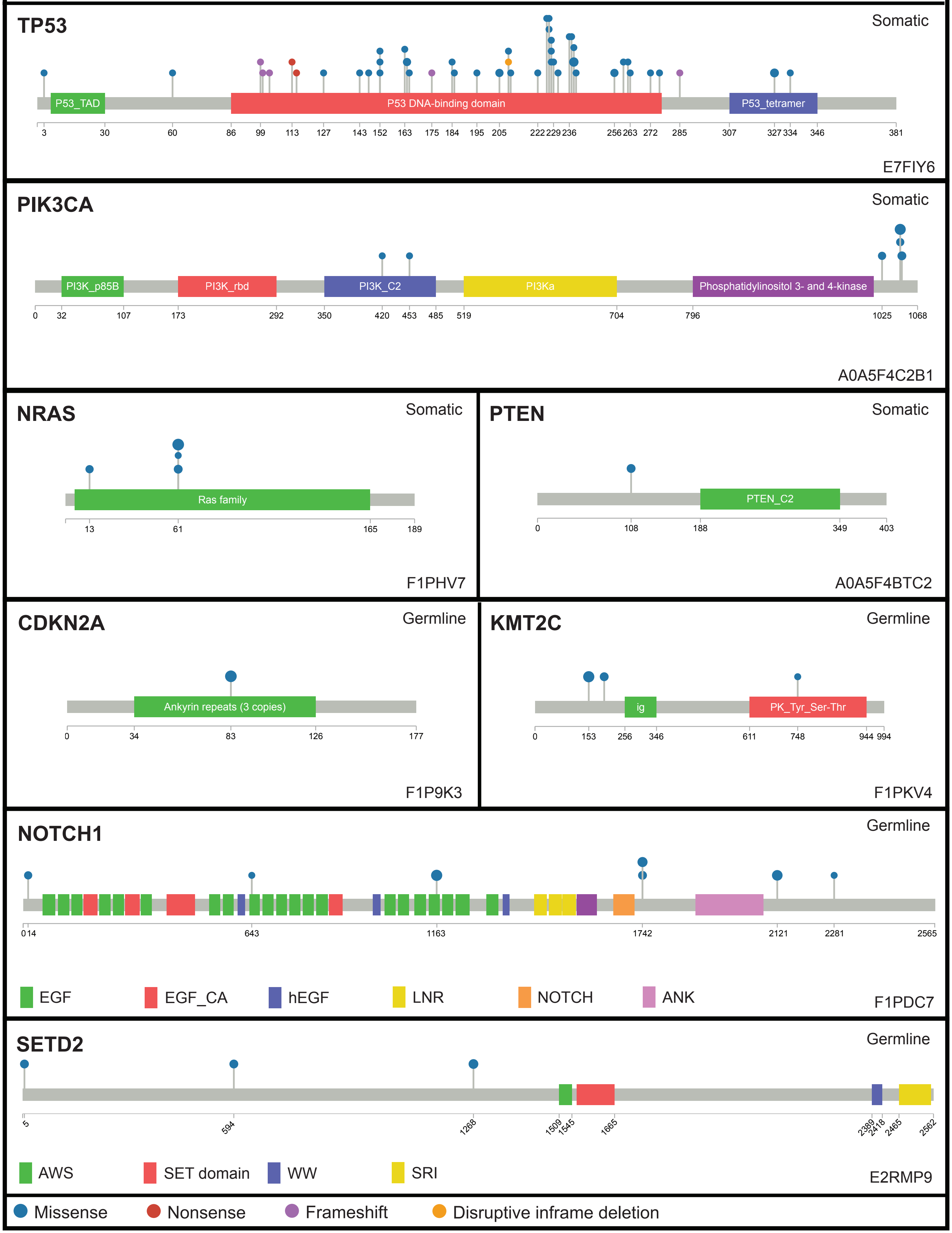

### Supplementary Figure 4

**A**

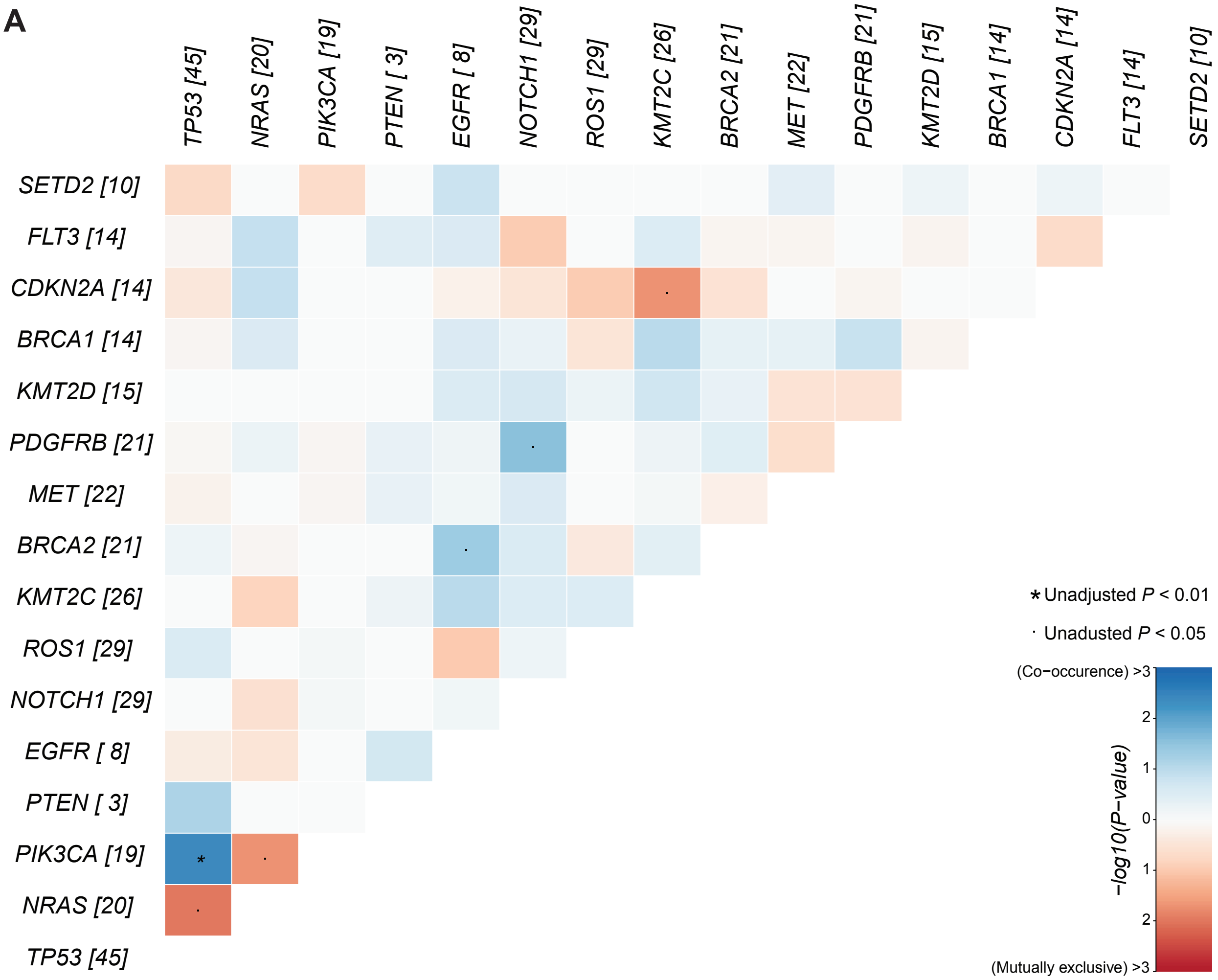


**B**

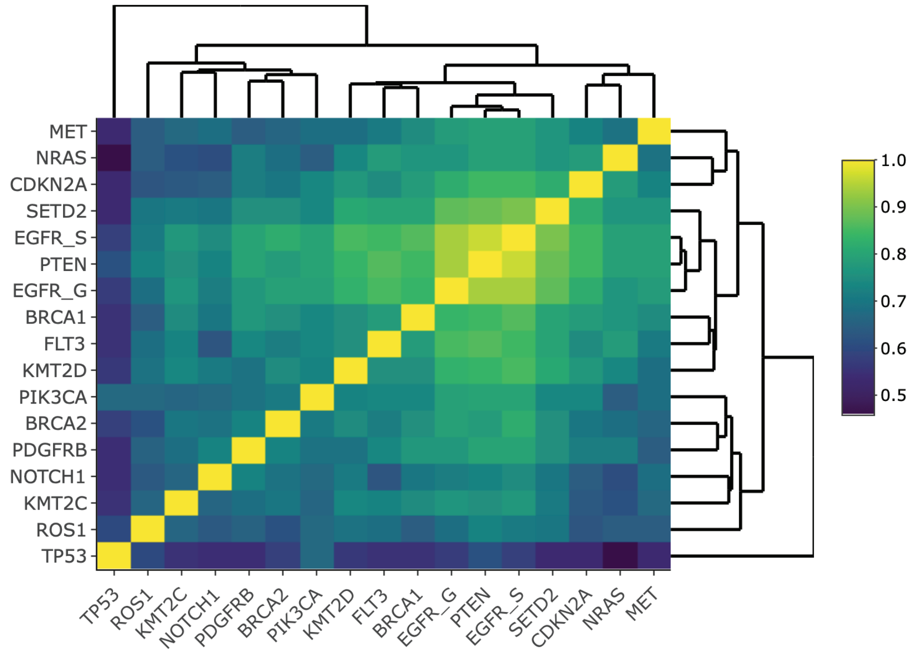

#### Supplementary Figure 5

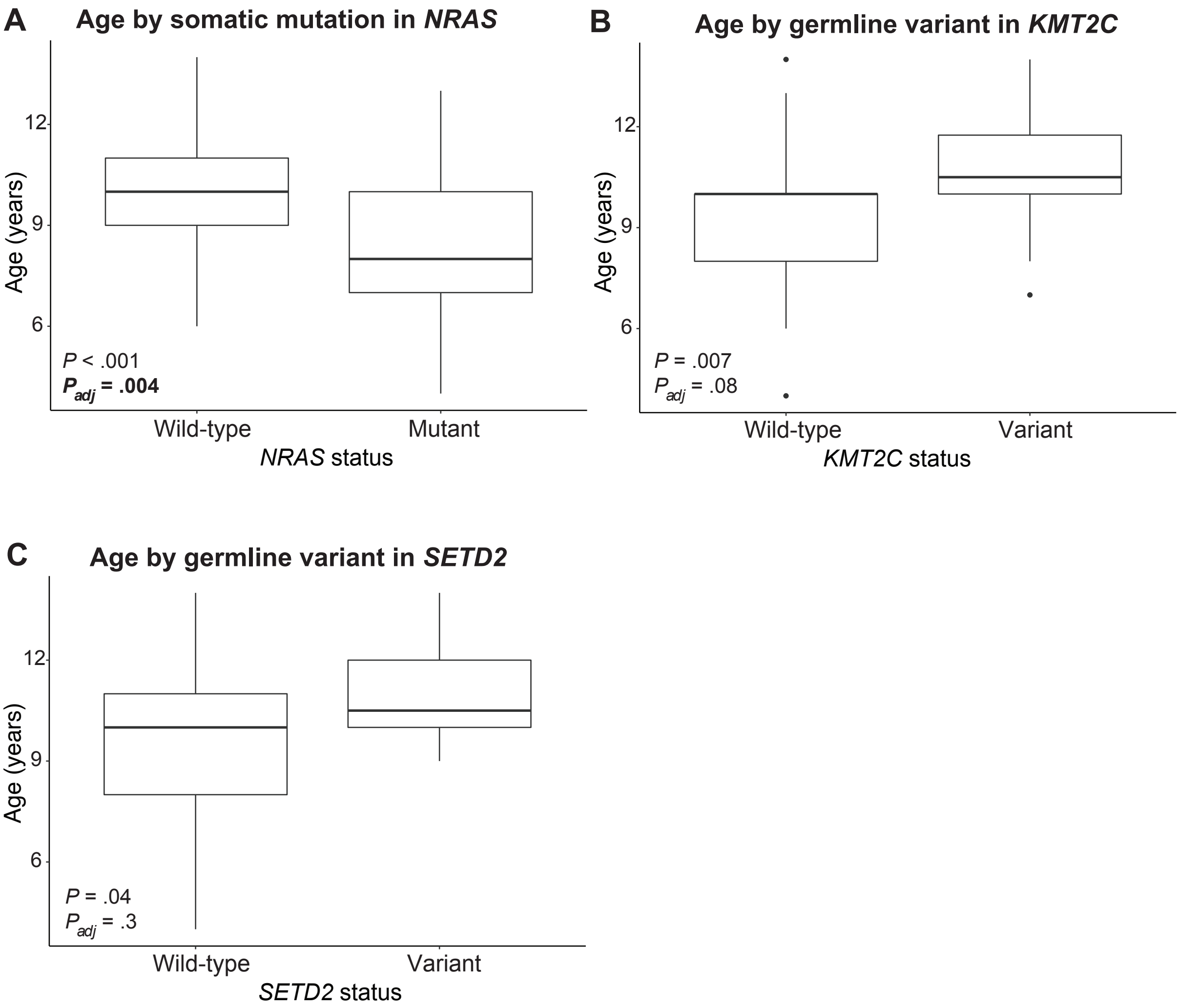

### Supplementary Figure 6

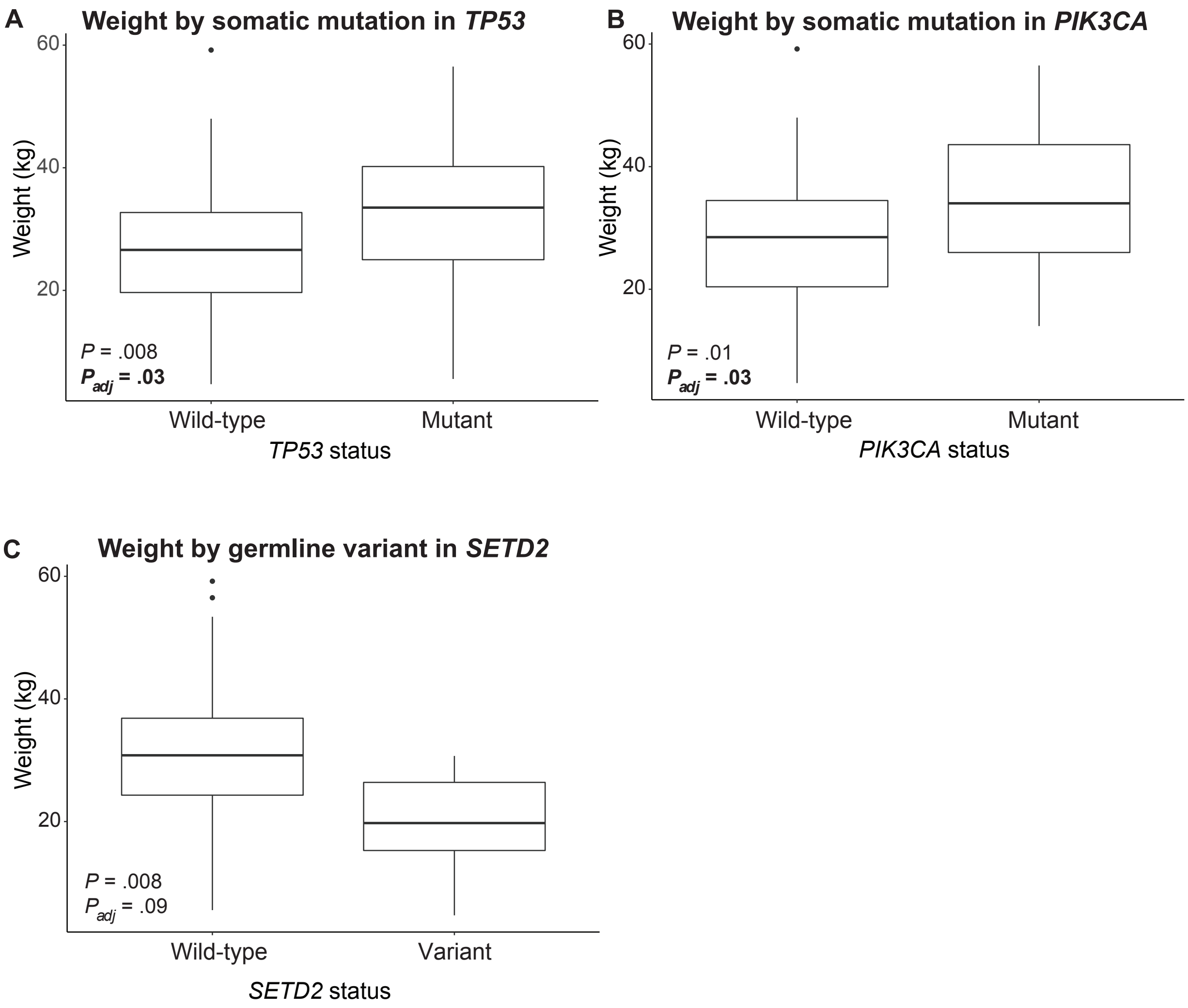

#### Supplementary Figure 7

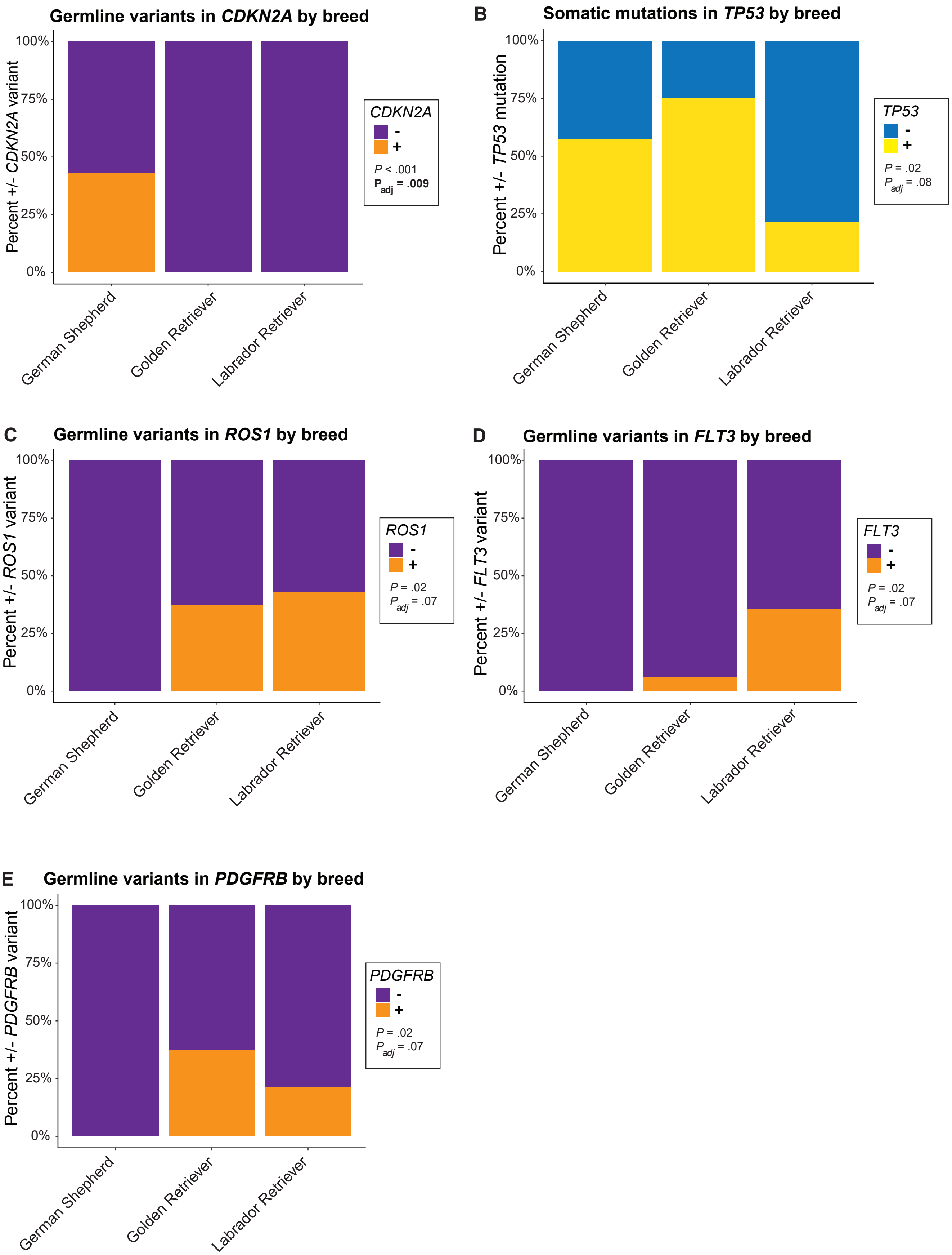
